# New, easy, quick and efficient DNA replication timing analysis by high-throughput approaches

**DOI:** 10.1101/858803

**Authors:** Djihad Hadjadj, Thomas Denecker, Eva Guérin, Su-Jung Kim, Fabien Fauchereau, Giuseppe Baldacci, Chrystelle Maric, Jean-Charles Cadoret

## Abstract

DNA replication must be faithful and follow a well-defined spatio-temporal program closely linked to transcriptional activity, epigenomic marks, intra-nuclear structures, mutation rate and cell fate determination. Among the readouts of the DNA replication spatio-temporal program, replication timing (RT) analyses require complex, precise and time-consuming experimental procedures, and the study of large-size computer files. We improved the RT protocol to speed it up and increase its quality and reproducibility. Also, we partly automated the RT protocol and developed a user-friendly software: the START-R suite (*<u>S</u>*imple *<u>T</u>*ool for the *<u>A</u>*nalysis of the *<u>R</u>*eplication *<u>T</u>*iming based on *<u>R</u>*). START-R suite is an open source web application using an R script and an HTML interface to analyze DNA replication timing in a given cell line with microarray or deep-sequencing results. This novel approach can be used by every biologist without requiring specific knowledge in bioinformatics. It also reduces the time required for generating and analyzing simultaneously data from several samples. START-R suite detects constant timing regions (CTR) but also, and this is a novelty, it identifies temporal transition regions (TTR) and detects significant differences between two experimental conditions. The informatic global analysis requires less than 10 minutes.

## Introduction

DNA replication is a highly regulated process involved in the maintenance of genome stability (Hanahan and Weinberg, 2011; Macheret and Halazonetis, 2015; Técher et al., 2017). Its accuracy relies partly on a spatio-temporal program that regulates timing and location of origin firing (Dileep et al., 2015; Rivera-Mulia and Gilbert, 2016). Based on this program, replication is organized into large-scale domains that replicate at different times in S phase (Ryba et al., 2010; Cornacchia et al., 2012). During the last decade, different groups including our laboratory, showed that the replication-timing program (RT) is finely tuned and maintained from an S phase to the following one (Hadjadj et al., 2016; Brustel et al., 2017; Almeida et al., 2018). It also appeared that this program is modified during cell differentiation (Hiratani et al., 2010; Gilbert 2012; Hadjadj et al., 2016). However, it remains unclear how this program is established and maintained. Protocols developed to study the RT in specific cell lines have been established in different labs (Hansen et al., 2010; Ryba et al., 2011; Dileep et al., 2012; Marchal et al., 2018). Differences between RT protocols may produce different results, sometimes devoid of biological relevance. DNA-Immunoprecipitation (DNA-IP) is a critical step of the RT protocol. We optimized duration and reproducibility of this step by using the SX-8G IP-Star® Compact Automated System (Diagenode®). Thus, it now lasts only 1 day (instead of 2-3 days before) and produces highly reproducible results regardless of the experimenter. In order to make the analysis of experimental results more accurate and reproducible, we also implemented two web-based softwares: START-R Analyzer and START-R Viewer, showing user-friendly interfaces (HTML and simple-click controls) that can be used by any biologist. The START-R Analyzer was initially based on a script developed in 2011 by David Gilbert’s laboratory (Ryba et al., 2011) which is not anymore currently working as it is, due to different software updates. The script was improved by implementing new tools for the detection of temporal transition regions (TTR) and for the fast identification of differential results between two experiments. These softwares are available online on our GitHub group website (https://github.com/thomasdenecker/START-R), they are free and each developer can improve them according to specific needs. We validated the START-R suite with Drosophila, zebrafish, mouse and human RT data obtained with microarrays or high-throughput sequencing. Using this automated DNA-IP protocol followed by analysis with the START-R Suite, it becomes easier for a large number of laboratories to carry out studies on the RT, thus opening up to new research perspectives.

## Results

### An improved RT protocol using the IP-Star robot

The protocol developed to analyze genome-wide replication-timing program (RT) in mammalian cells lasts 3 weeks (Ryba et al., 2011). It includes pulse-labeling of cells with nucleotide analog 5-bromo-2-deoxyuridine (BrdU) followed by flow cytometry cell sorting (FACS) of labeled cells into two S-phase fractions. Then, immunoprecipitation (IP) targeting the BrdU-labeled DNA is performed. IP is a time-consuming step and needs to be carefully monitored in order to get precise and specific signals. Thus, we optimized the BrdU IP protocol by introducing an automated step using the SX-8G IP-Star® Compact Automated System. It is now possible to simultaneously perform IP of 16 samples that correspond to 8 early and 8 late fractions, which means 8 RT experiments. The automated system allows to perform DNA-IP overnight. Its high standardization improves the reproducibility of RT profiles from one experiment to another, whoever the experimenter. To prove this point, RT analyses were performed by four different experimenters with four independent RKO cell line cultures; two experimenters performed handmade DNA-IPs independently, while two others independently used the IP-Star robot. Then, we compared the percentage of differences between each handmade experiment and each “IP-Star” experiment. We found a difference of 4.25% between both handmade experiments while only a difference of 0.11% was observed between each independent experiment performed with the “IP-Star” robot (Supplemental Fig. S2 and Supplemental Table S1).

Once immunoprecipitated, newly synthesized DNA is amplified with a SeqPlex™ enhanced DNA Amplification kit designed for microarray or deep-sequencing experiments. For microarray experiments, labeled DNA is hybridized to a whole genome comparative hybridization microarray (CGH microarray, 180,000 probes, one every 13Kb). We showed that a microarray with only 60,000 probes is not sufficient to produce a detailed RT profile (Supplemental Fig. S3). After scanning, the generated picture is analyzed through the feature extraction software (Feature Extraction 9.1 Agilent) that measures Cy3 and Cy5 intensity values for each of the 180,000 probes of the microarray. This generates a table containing the measures of processed signals that are used by START-R. “SystematicName”, “gProcessedSignal and rProcessedSignal” columns are the ones used by default by START-R Analyzer to generate the whole-genome RT profile and its analysis. Other column names could also be used but experimenters have to be sure that such names are well matched when the file is downloaded to START-R Analyzer.

### The Repli-chip protocol leads to results similar to the “6 fractions Repli-Seq” protocol

To test the accuracy of our Repli-chip approach, we compared it with the previously used “6 fractions-Replication-sequencing (Repli-seq)” method (Hansen et al., 2010). Both approaches were applied to the same K562 cell line. We retrieved these Repli-seq data (6 fractions corresponding to G1, S1, S2, S3, S4 and G2) from the UCSC genome browser website (Kent et al., 2002).

Initially, we chose the position of cell sorting windows in S phase in our Repli-chip experiment on the basis of previous validated RT protocols (blue lines, Fig. 1A). When S1(Early) and S2 (Late) fractions are limited only to the S-Phase during Repli-chip experiments (blue lines, Fig. 1A), some regions finally appeared as replicating in the middle of S phase (blue smooth RT profile, Fig. 1C), whilst they had been shown to replicate very late in the “6 fractions Repli-seq” experiment (Fig. 1B). This could be explained by the low amounts of nascent DNA fragments present in the Early and Late sorted fractions restricted to S phase. The analysis of this Early/Late ratio results in an artefactual mid S phase replicating domain (blue smooth RT profile, Fig. 1C). Thus, these regions appear to replicate in the middle of S phase while corresponding to domains replicated during the very late S-phase and the beginning of G2.

**Figure 1.**
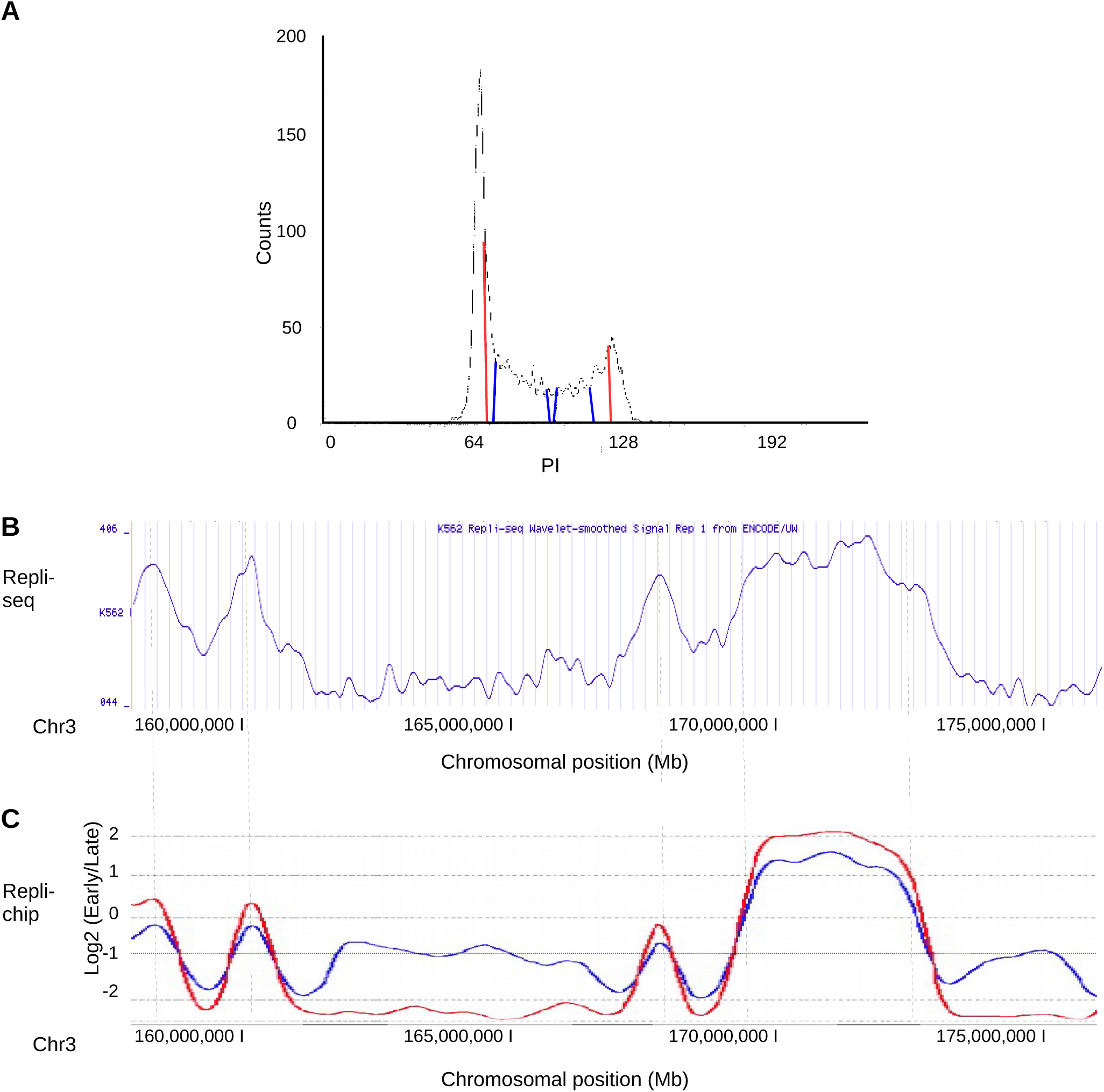
Repli-chip and Repli-seq protocols lead to similar results. **A)** Cell cycle profile of K562 cells after propidium iodide labeling. The blue lines indicate windows used to sort cells in the first (S1) and the second part of S phase (S2). The red lines indicate the new wider delimitations of Early and Late sorting windows. **B)** Replication profile of K562 cell line obtained with Repli-seq approach. **C)** Replication profiles of K562 cell line obtained with our Repli-chip protocol. The blue line depicts the replication profile obtained after using the sorting windows limited to the S phase. The red line depicts replication profiles obtained using wider sorting windows overlapping slightly with G1 (on the left) and G2 (on the right) phases of cell cycle (as shown in A). The x-axis displays chromosomal positions (Mb) and the y-axis log2 (Early/Late) intensities. Dashed vertical lines show common RT regions between the B and C profiles.

To fix this problem, we have expanded the cell sorting windows to the end of G1 for the Early fraction, and into the beginning of G2 for the Late fractions (red lines, Fig. 1A). Using these cell-sorting parameters RT profiles are highly similar to the “6 fractions-Repli-seq” experiments despite the gap left between S1 and S2 fractions to avoid cross-contaminations between both fractions, as shown for chromosome 19 and all other chromosomes (red smooth RT profile, Figs. 1B, 1C and Supplemental Fig. S4, respectively). Therefore, an accurate Repli-chip protocol with correct cell-sorting parameters for two fractions provides similar results to those produced by the “6 fractions Repli-Seq” approach, in a less expensive and less time-consuming way.

### START-R suite for automation of RT analysis allows robust statistical analysis with a user-friendly interface

We developed a software suite starting from a script created by David Gilbert’s group in 2011 (Ryba et al., 2011), that we improved and updated with different current versions of tools and algorithms. We also implemented new functions as TTR detection and differential analysis. The START-R suite, which stands for *<u>S</u>*imple *<u>T</u>*ool for the *<u>A</u>*nalysis of the *<u>R</u>*eplication *<u>T</u>*iming based on *<u>R</u>*, is implemented into an HTML interface for more efficient and easier installation, use and sharing by biologists. START-R is built-in with Docker that packages START-R into a virtual container (Supplemental Fig. S1). Thus, START-R can be easily deployed at a personal computer or on a server, and can run independently of any library updating. This was not the case in the script developed in 2011 (Ryba et al., 2011), which makes that script much less easy to use (Supplemental Fig. S1). As indicated by its acronym, START-R has the strength of being based on a statistical approach using R, allowing researchers non-initiated in R programming to analyze their data. While the usability of the program would tend to limit its adaptability, START-R provides as many parameters as possible for a comprehensive analysis of the RT program (Supplemental Fig. S5A to 5K). Furthermore, we added new scaling, normalization and smoothing methods (Supplemental Figs. S6, S7) and also novel statistical approaches to detect differences between two samples (Supplemental Fig. S8, S9). A classical differential analysis performed with START-R takes only 5-6 min (compared to several hours without using START-R), with a personal computer containing an intel® Xeon(R) CPU E5-1620-3.60GHz × 8 core and 32GB of memory with the 18.04 Ubuntu version. START-R Analyzer runs by default with the hg18 human genomic annotations including the position of the centromeres. It can also be used without the centromere positions or with the centromere positions uploaded in a corresponding chromosome coordinates file for all organisms and all annotations (>hg18 for human genome). This flexibility is one of the new aspects of the START-R suite that allows to analyze RT program in every organisms (Supplemental Fig. S5B).

### A large panel of new settings and tools for RT analysis

In our method, we based our script on four major steps: <u>normalisation</u> (between Early and Late fractions, between two replicates, and between two independent experiments, Supplemental Fig. S6), <u>smoothing</u> (LOESS, Simple, Weighted, Modified, Triangular, Exponential and Running methods including limma, Ritchie et al., 2015, Supplemental Fig. S7), <u>identification</u> of transition timing regions (TTRs), and <u>segmentation</u>. The originality of our approach is to first detect TTRs in order to better identify Constant Timing Regions (CTRs, Supplemental Figs. S8A, S8B). The identification of TTRs is based on their intrinsic properties: regions that include more than three consecutive probes with significantly different Early/Late intensity log ratios are considered as TTRs (Supplemental Fig. S8A). The statistical significance of differences between probes is calculated by the outlier boxplot method (Supplemental Fig. S9, Krzywinski and Altman, 2013). Following TTRs detection, START-R Analyzer localizes CTRs: TTRs are subtracted from the genome (Supplemental Fig. S8B) and the remaining regions are considered as CTRs. If TTRs were not excluded initially, the CTRs would overlap the adjacent TTRs (Supplemental Fig. S8B). This overlap would result in a less precise segmentation because the algorithm would take into account the probes positioned in adjacent TTRs for calculating the segment value. After subtraction of TTRs, START-R Analyzer scans the remaining regions through a sliding window-based algorithm. It will potentially divide these regions into different CTRs if the standard deviation values of intensities reach a chosen threshold. At the end of these steps, START-R Analyzer automatically generates a BED file for CTRs and TTRs making easier further bioinformatics analyses and the display of the RT domains via a genome browser (Supplemental Fig. S5F). It also produces descriptive statistics of the normalization step and RT statistical elements for each chromosome (domain size distribution, summary of segmentation process). Finally, a codebook is generated to ensure the traceability of options chosen for each analysis. The user-friendly interface facilitates the choice among the different analysis parameters (Supplemental Figs. S5A to S5K).

We added a step allowing the differential analysis of RT programs from two experiments. Thus, we can now compare RT profiles obtained in different conditions and/or with different cell lines to identify loci and elements that can modify the RT program. Our differential analysis includes three different methods of comparison: the Mean method, the Euclidean method and the Segment comparison method.

The Mean method compares the means of log ratio intensities (Early/Late) obtained in two different experiments. Mean is calculated for a sliding window of 30 successive probes corresponding to a 300 kb genomic domain, which is consistent with the size of replication-timing domains already described (Rivera-Mulia and Gilbert, 2016). The overlapping parameter, defining the number of probes overlapping successive windows was initially set to 15. After the calculation of nominal and adjusted p-values (t-tests for mean comparison), the user can choose the p-value thresholds to distinguish significant differences between two conditions. The p-value adjustment method is chosen among a list of classical procedures (such as Bonferroni, Holm and Benjamini and Hochberg methods).

The Euclidean method computes the squared differences of the log ratio intensities (Early/Late) for the same probes in two different experiments. The squared differences of every probes are plotted in a boxplot and the outliers are considered as significantly different following the threshold chosen by users (Supplemental Fig. S9). Regions of differential intensities are defined as more than three consecutive outlier probes.

The segment approach uses the TTR and CTR values generated by START-R in the TTR and segmentation steps. The user can choose any parameter as this approach employs an empiric method. First, the method compares the presence or the absence of TTRs in two experiments (Supplemental Fig. S8C). If TTRs are detected in both experiments, the segment approach compares their slopes(Supplemental Fig. S8D). Finally, means of probe intensities are compared between CTRs common to two different experiments, by t-test. The threshold for the t-test significance is calculated according to the following procedure: two empiric CTRs are generated by picking randomly intensity values in both CTR for each experiment and a p-value is calculated by t-test. This permutation process is repeated 10.000 times and results to the estimation of an empiric p-value that is the threshold p-value for CTR comparison. The last major implementation is START-R Viewer (Supplemental Fig. S5K). This web-based interface allows the visualization of the RT profile generated by START-R Analyzer in dynamic charts obtained with the Plotly library (Sievert et al., 2017). It creates figures from a specific file generated by START-R Analyzer, integrating all the analyses performed by START-R analyzer that another genome browser cannot display optimally. One can easily identify CTRs, TTRs (Fig. 2A) and significantly advanced or delayed regions (Fig. 2B). The user can also choose color options for the RT display and take automatically a screenshot of the RT profile to create a figure. We therefore developed a genome browser to optimally display the maximum of resources generated by START-R Analyzer.

**Figure 2.**
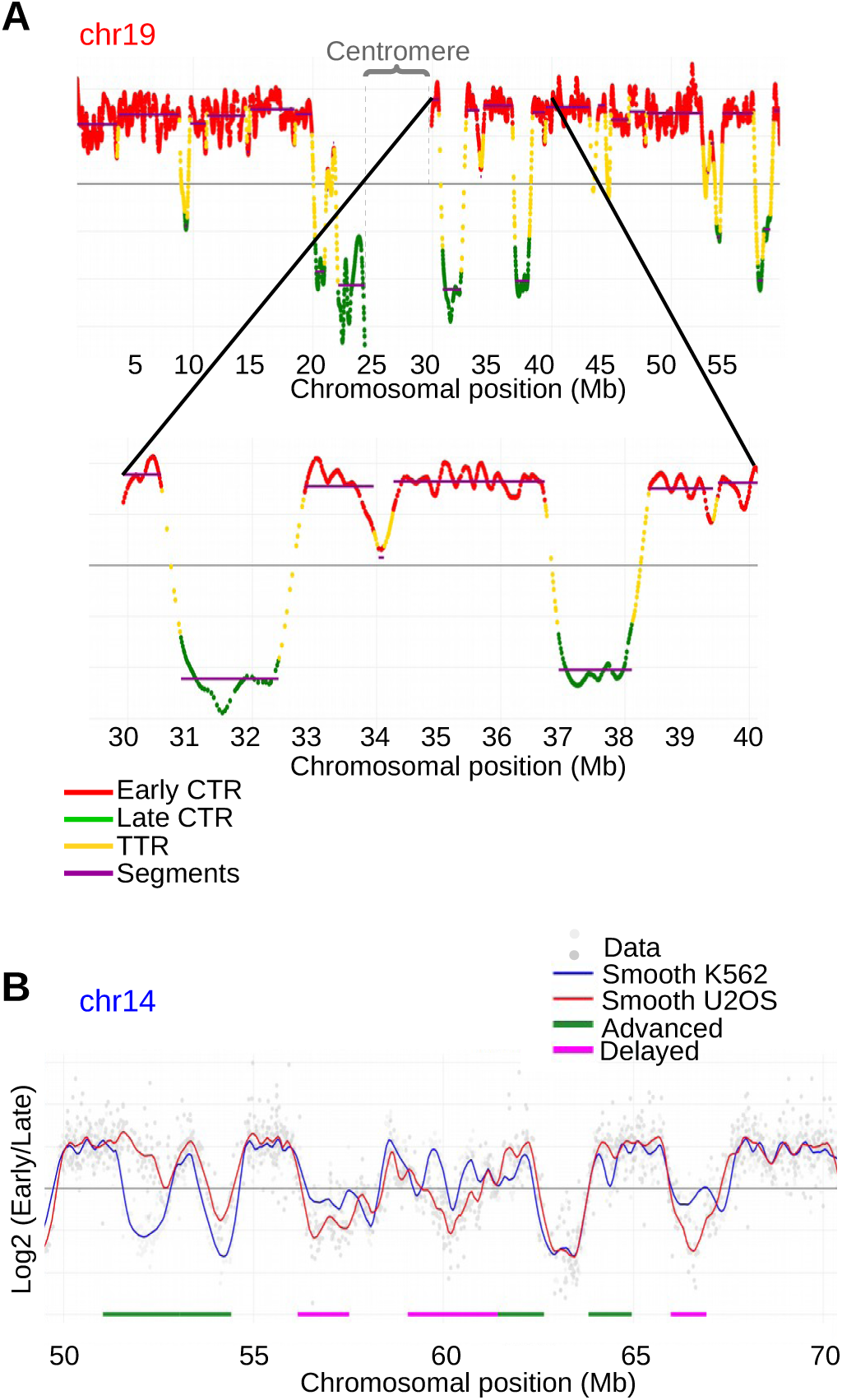
Examples of data generated with our RT protocol visualized by START-R-Viewer. A) START-R Viewer allows visualizing RT data with many features. The top panel displays the distribution of early and late constant timing regions (CTR, in red and green, respectively) and of transition timing regions (TTR, in yellow) on a portion of human chromosome 19. Segments corresponding to regions of constant timing are shown in purple. Chromosome 19 centromere is indicated by grey dashed lines and a curly bracket. The bottom panel displays a zoom of a smaller region of chromosome 19 where timing profile can be seen through the zoom option of START-R-Viewer. B) Differential analyses are done on a portion of human chromosome 14 comparing RT profiles of two cell lines: K562 in blue and U2OS in red. Advanced (green) and Delayed (pink) regions are identified with START-R Analyzer using the mean comparison analysis with the Holm’s p-value correction and a limit corrected p-value of 0.05. Light grey and grey spots indicate data from both RT experiments.

### How to choose adapted thresholds for differential analyses?

As described above, START-R Analyzer proposes different parameters depending on the chosen method. The central point for users is to select the correct method for the differential analysis and to choose the adapted parameters for this process. To test the impact of parameters on the detection of differences between two conditions, we compared RT data from the K562 and U2OS cell lines. We used the same normalization, smoothing, TTR and segment detection methods for both cell lines. We tested the Mean, Segment and Euclidean methods for differential analyses (Fig. 3A, 3B and 3C). Only Mean and Euclidean methods offer the possibility to change threshold parameters. The segment method does not use p-value thresholds, or thresholds, since it is based on empirical parameters.

**Figure 3.**
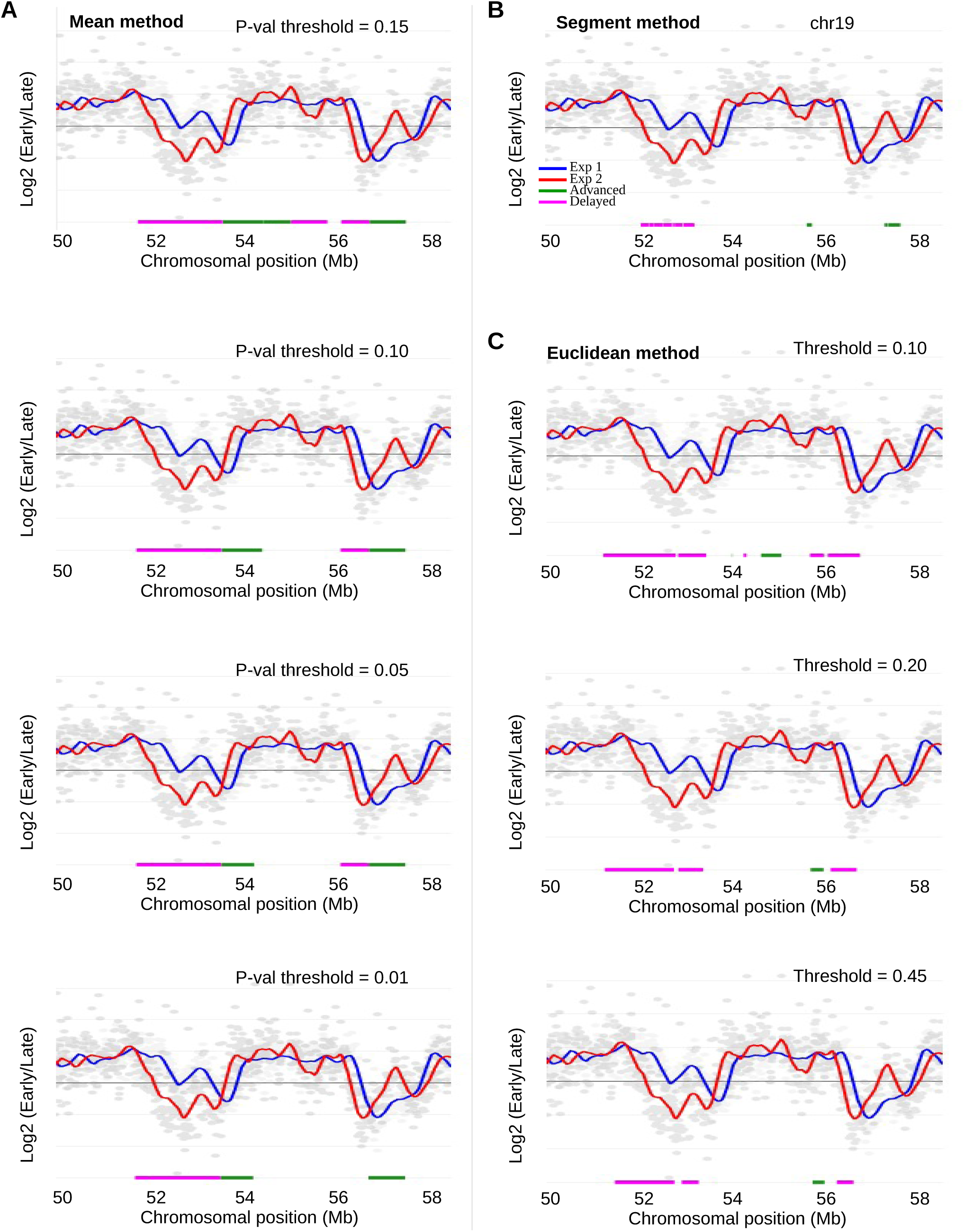
Comparison of differential analyses of RT profiles with START-R Analyzer using the Mean, Segment and Euclidean methods. **A)** Differential analyses allow the comparison of RT profiles of a portion of chromosome 19 for two cell lines, K562 (blue) and U2OS (red). Light grey and grey spots indicate data from both RT experiments. The left panels show the identification of Advanced (green) and Delayed (pink) regions using the Mean method with a corrected p-value threshold ranging from 0.15 to more stringent p-values of 0.10, 0.05 and 0.01, respectively. **B)** Differential analysis of the same chromosomal region using the Segment method based on empirical parameters. **C)** Differential analyses of the same chromosomal region with the Euclidean method with a threshold varying from 0.10 to 0.45.

To further explore the relationship between the p-value threshold or threshold and the detection sensitivity of true RT changes with the Mean and Euclidean methods, we examined the significant timing domain changes when the p-value threshold, or the threshold, increase (Fig. 3A and 3C). We used graphs depicting the significant timing change coverage related with the p-value threshold, or threshold (Figs. 4A and 4B and Supplemental Fig. S10). We defined an optimized p-value threshold, or threshold, as the parameter at which the gain of additional RT change coverage is minimal. For each method, we draw the chord of the curve (Figs. 4A and 4B and Supplemental Fig. S10). The perpendicular and longer segment between the chord and the curve was defined, indicating the optimized parameter (Supplemental Fig. S10). These tests showed optimized p-value threshold and threshold for the Mean and Euclidean methods of 0.025 and 0.192, respectively (Supplemental Fig. S10A and S10B). Using these parameters, we found 713 RT changing regions with the Mean method, 1798 with the Euclidean method, and 3397 with the Segment method (Fig. 4C). Each method shows its particularities, however, 659 common RT changing regions were detected by the three methods. While the number of regions with RT changes is different for each method, the global genome coverage is the same. As START-R Analyzer works quickly, we propose that experimenters use the same procedure in order to determine their own optimized parameters suitable for their experiments.

**Figure 4.**
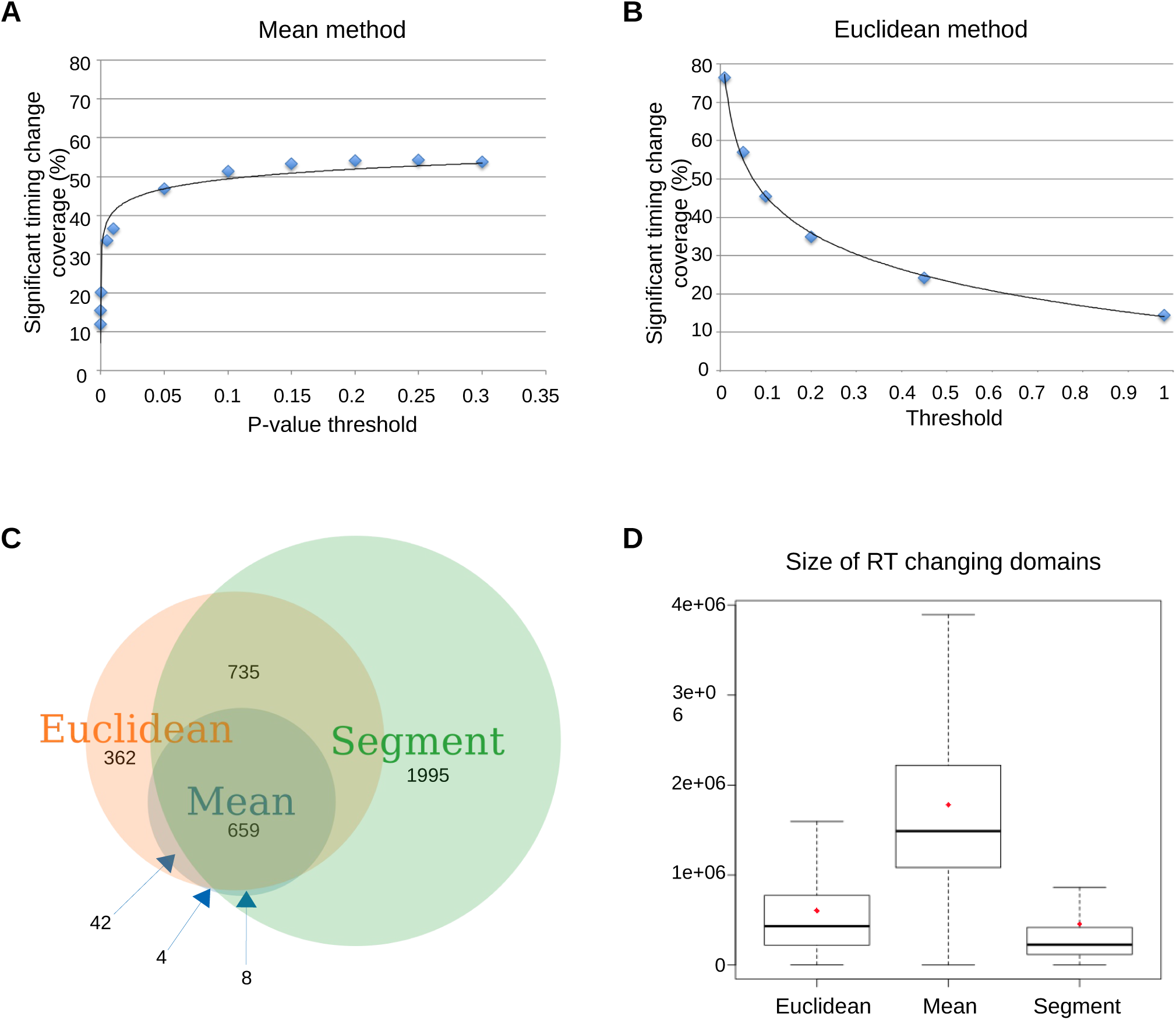
Comparison of the different analysis methods. **A) and B)** Graphs are depicting the increase in percentage of the significant timing change coverage with an increased p-value threshold (A, Mean method) or an increased threshold (B, Euclidean method). **C)** Venn diagram depicts the comparison between the 3 different methods (Mean, Segment and Euclidean) with optimized calculated p-value threshold (0.025) and threshold (0.192, see Supplemental Figure 9 for additional details) to detect common RT changes (advanced and delayed regions). We used the overlapping intervals option from the Intersect intervals of 2 datasets tools in Galaxy. **D)** Boxplots illustrate differences in the size of RT changing domains between Euclidean, Mean and Segment methods. For each category the mean value is indicated by a red diamond. The band at the middle of the box indicates the median value. The bottom and top of the box are the 25^th^ and 75^th^ percentiles.

Then, we compared the three methods with different combinations of parameters (Supplemental Fig. S11) in order to evaluate the capacity of each method to detect significant RT changes. As part of a comparison between the K562 and U2OS cell lines, the Segment method allows the detection of the highest number of RT changes (3397 regions with an average size of ≃ 450Mb). The Mean Method shows the lowest RT changes (around 710 regions with an average size of ≃ 1,800Mb), representing around 18-21% of RT changes detected by the Segment method (Supplemental Fig. S11). The Euclidean method reveals an intermediate number of RT changes (around 1932 regions with an average size of ≃ 607 Mb). Thus, each method shows a specific range concerning the length of regions with RT changes (Fig. 4D).

Nevertheless, 92 to 95% RT changes detected by the Mean method overlap those detected with the other two methods (Supplemental Fig. S11). 40-48% RT changes detected by the Euclidean method overlap those detected by the Segment method, and 29-40% overlap those detected by the Mean method (Supplemental Fig. S11). With the optimized parameters, only 0.5% of RT changes found by the Mean method are unique, compared to 20% for the Euclidean method and 58.7% for the Segment method (Fig. 4C). These observations show that a large part of RT changes, but not all, are detected by the Mean method. The Segment method appears to be more sensitive and able to detect more new regions that the other two methods. Each method has its own detection characteristics. Each user, depending on the asked biological questions, has to choose the most suitable one for the analysis. When the goal is to identify most regions with a real RT change, we recommend using the Mean method. When the goal is to find all regions with a RT change, we recommend using Segment or Euclidian method but with increased risks of obtaining false positives. Since the analyses with START-R are fast, we also recommend performing the analysis using the three methods and keeping the common results. Therefore, it is important that START-R Analyzer proposes these three tools.

### START-R analysis of replication-timing programs during differentiation in mouse: a new analysis of previous data

To validate our START-R based-approach without *a priori* consideration, we decided to reanalyze the data obtained by the Gilbert’s group concerning the changes of replication-timing program during cell differentiation in mouse (Hiratani et al., 2008). They found that 20% of replication domains change between the D3esc and D3npc9 cell lines. There are two types of changes: Early-to-Late (EtoL or delayed) and Late-to-Early (LtoE or advanced). Each modified timing region had a particular molecular signature: LtoE regions show a GC/LINE-1 density and gene coverage similar to constant early regions, while EtoL regions showed GC/LINE-1 density and gene coverage similar to constant late regions. We used the same raw data for START-R analysis. First, we converted the raw data with the convertPair.R script to be in the correct format for START-R Analyzer. Then, we used START-R Analyzer with the standard option: Loess Early/Late normalisation, scale inter-replica normalisation, inter-experiments standardization, Loess method for smoothing (span=300kb), 2.5 for SD difference between two segments, and Holm’s method with a p-value=0.05 for the differential analysis. With these parameters, 2,066 CTRs are detected in the genome and 910 regions show a different replication timing between D3esc and D3npc9 (Fig. 5A). Thanks to START-R Analyzer that automatically generates BED files, it is easy to import files into a GALAXY session (Afgan et al., 2018) in order to continue the molecular characterization and to generate complementary results. Advanced and delayed regions show the aforementioned specific molecular signatures for the GC/LINE-1 content and gene coverage (Fig. 5B; Hiratani et al., 2008). Unfortunately, we cannot compare the regions that we have identified with START-R with those previously discovered, since the article of Hiratani *et al* does not mention the genome coordinates. However, our results confirm that START-R Analyzer is robust, whatever the mammalian replication timing program studied and whatever the type of microarrays used (from Agilent or Affimetrix or Nimblegen). In fact, we detected identical molecular signatures in the regions showing RT changes.

**Figure 5.**
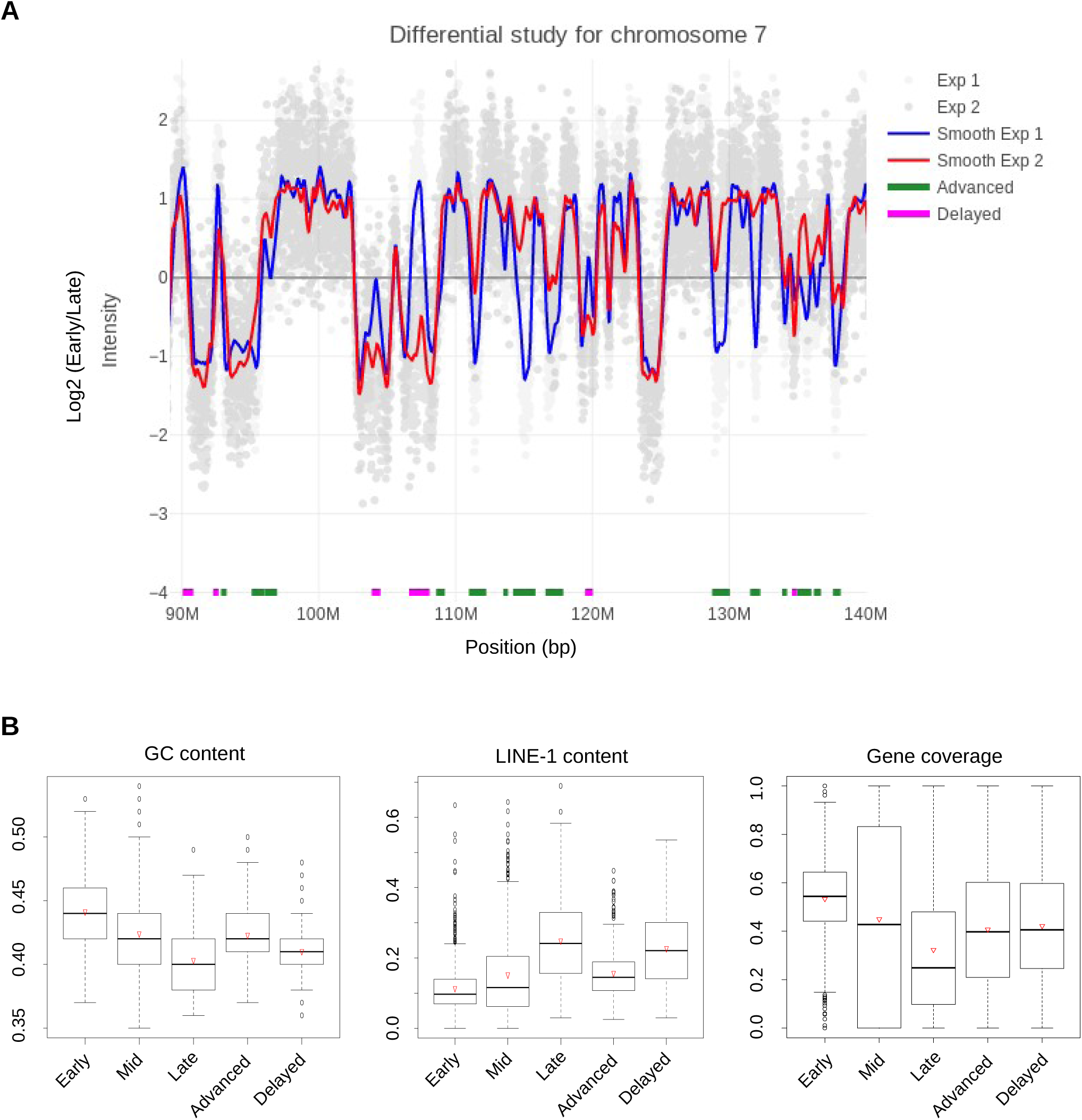
Genomic characteristics of regions harboring different replication timing programs. **A)** START-R Differential analysis of RT profiles is shown for a portion of chromosome 7 in mouse D3esc (blue) and D3npc9 (red) cells. Light grey and grey spots indicate data from both RT experiments. **B)** Boxplots illustrate differences in GC content, LINE-1 content and gene coverage between Early, Mid and Late replicating regions. The two other categories show the characteristics of Advanced and Delayed regions. For each category, the mean value is indicated by an open red triangle. The band at the middle of the box indicates the median value. The bottom and top of the box are the 25^th^ and 75^th^ percentiles. Bottom and top whiskers represent the limits with exclusion of outliers (open circles).

### Validation of START-R with Early-Late Repli-seq data from mouse

Many data of replication-timing program can be obtained with Repli-seq experiments, but their analysis is time-consuming and often require bioinformatics skills. We analyzed the Early/Late repli-seq data from Marchal and co-workers (Marchal et al., 2018). With different genomic tools (using GALAXY in our case), we obtained the alignment of reads and a BAM coverage file essential for the integration in the START-R pipeline. We specifically developed a supplemental script to convert the BAM coverage file to a log Early/Late file (convert_bamcoverage_file.R) to be sure that the integration into the START-R pipeline was correct. Then, we compared the RT smooth profile from E/L Repli-seq with similar data obtained with microarrays (Fig. 6A). The profiles are almost identical, exactly as described by Marchal and co-workers (Marchal *et al*., 2018). Thus, START-R Analyzer and Viewer can be easily used to analyze E/L Repli-seq data, showing its versatility and its simplicity of use.

**Figure 6.**
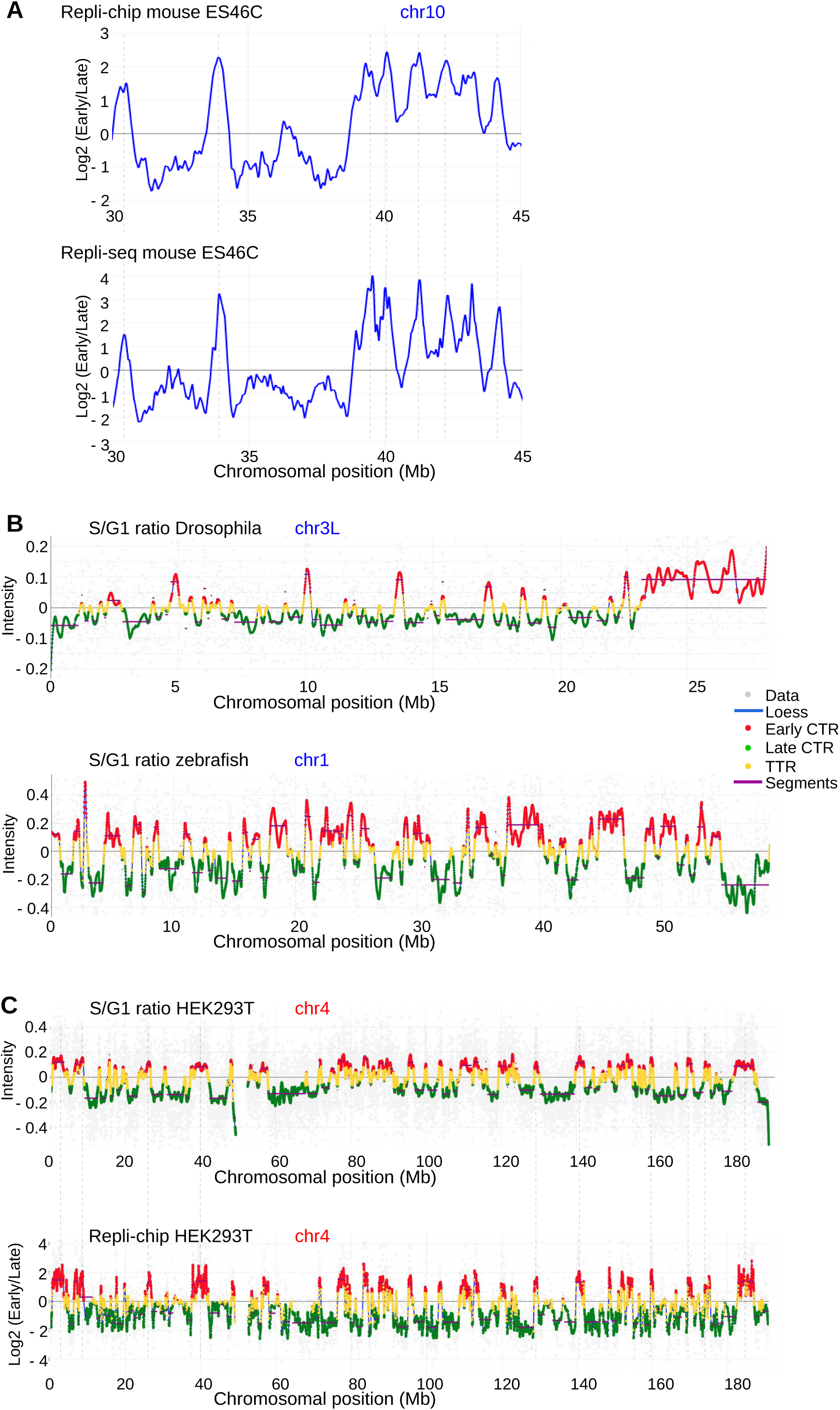
The START-R suite allows analysis and visualisation of both Repli-chip and Repli-seq data from different model systems. **A)** RT profiles of a portion of mouse chromosome 10 from ES46C cell line are generated using Repli-chip (top panel) and Repli-seq data (bottom panel) with START-R software. Dashed vertical lines show common RT regions between both profiles. **B)** RT profiles obtained by S/G1 ratios are shown for the left part of Drosophila chromosome 3 (3L) and for zebrafish chromosome 1 (blue lines). The profiles display distribution of early and late CTRs, in red and green, respectively and of TTRs, in yellow. Segments corresponding to regions of constant timing are shown in purple. Grey spots indicate data from RT experiments. **C)** RT profiles of human HEK293T chromosome 4 are generated using S/G1 ratio and Repli-chip data. The empty space inside the RT profiles represent the centomere region.

### Validation of START-R with S-G1 Repli-seq data from Drosophila, zebrafish and human

Other laboratories use the ratio of DNA content between G1 and S phases to analyze the RT program. We wanted to know if START-R suite can run the correct analyses with this type of data and also with other organisms than mouse and human. We performed exactly the same pipeline used for early-late Repli-seq data described above for Drosophila, zebrafish and human S/G1 data (Armstrong et al., 2018; Siefert et al., 2017; Massey et al., 2019), in order to be sure that the integration into the START-R pipeline was correct. Then and as expected, START-R can be run with S/G1 log ratio data for Drosophila, zebrafish and human (Fig. 6B and 6C). We observed similar profiles as the ones already observed for these different organisms. However, our analysis of HEK293T RT changes with the Repli-chip method gave higher differences between distant extreme values, which optimized the detection of RT changes.

## Discussion

In this study, we show a new automated protocol for generating and analyzing RT profiles in human and mouse genomes. This approach relies on both the automation of the IP step and on new web-based softwares, START-R Analyzer and Viewer (see graphical abstract in Supplemental Material). The IP-Star® robot reduces the length of the IP step from 3 days to an overnight experiment allowing the user to test 16 samples at the same time. This protocol is very interesting because samples could then be treated either by Repli-seq (2-fractions) or by microarrays. In addition, since the main experiment is carried out by a robot, there are few differences due to different experimenters. As demonstrated by our results, the degree of reproducibility of experiments using the IP-Star® robot is very high (Supplemental Fig. S2). In addition, the choice of the cell-sorting window is primordial. The window extension overlapping G1 and G2/M (Fig. 1A) gives exactly the same profile obtained with the “6-fractions Repli-seq” method (Fig. 1B and 1C), in shorter time and reduced financial costs. The “6-fractions Repli-seq” approach, which is long and expensive, can discourage a number of labs. This approach makes these experiments much more affordable with the same level of precision.

The START-R suite facilitates the analysis by making it more accessible to non-bioinformatician researchers. User-friendly interfaces integrate all used steps to generate RT profiles (Fig. 2) and users can choose different parameters at every step. Compared to the previous, no longer in use method (Ryba et al., 2011), START-R Analyzer detects TTR regions and better refines and improves the CTRs detection. In addition, START-R Analyzer contains new calculation methods for the identification of differences between two conditions or two cell lines (Fig. 3). This flexibility gives the users the opportunity to choose the differential analysis method and different parameters according to their questions (Fig. 4, Supplemental Figs. S10 and S11). START-R also processes data from different organisms (Figs. 5 and 6) and those obtained with different methods, such as two fractions Repli-seq, two fractions Repli-chip and S/G1 fractions data. It also automatically generates files with different output formats essential for further molecular characterizations and compatible for classical bioinformatic tools and/or for GALAXY genomic tools. START-R Analyzer is not exclusively developed for mammalian genome as we also generated RT analyses for Drosophila and zebrafish genomes (Fig. 6).

START-R Viewer produces a nice interface to visualize all the data generated by START-R Analyzer. It facilitates the analysis of RT. In addition, it makes easier the navigation along the genome to take screenshots suitable for future figures (Supplemental Fig. S5K).

START-R Analyzer and START-R viewer freewares are available on GitHub and their source codes are open to anyone who wants to improve and to integrate them into a personal computer or server, whatever the operating system. Finally, the START-R suite is very flexible since it can use data from different microarray platforms, from Repli-seq experiments (with 2 fractions or S/G1 fractions), and from different organisms (Figs. 2, 3, 5 and 6). In conclusion, it is now possible for any biologist or laboratory to readily explore new or previous replication timing data simply and quickly. Thus, a large number of laboratories can today use our approach and our softwares to find out if their experimental conditions are affecting the replication timing process or are correlated with other molecular mechanisms. START-R also allows to determine what parts of the genome are impacted and to characterize further those loci. Thanks to the accessibility of our approaches and softwares, their speed and efficiency, new research perspectives can be efficiently envisaged.

## Methods

### Experimental procedures to obtain early and late DNA replicating fractions

#### BrdU incorporation and cell fixation

3.10^7^ exponentially growing mammalian cells were incubated with 0.5 mM BrdU (Abcam, #142567), protected from light, at 37°C for 90 minutes. Cells fixed in 75% final cold EtOH can be stored at −20°C.

#### Cell sorting

10^7^ BrdU labeled cells were incubated with 80 μg/mL Propidium Iodide (Invitrogen, P3566) and with 0,4 mg/ml RNaseA (Roche, 10109169001) for 1h at room temperature with orbital shaking at 180 rpm. 10^5^ cells were sorted in early and late S phase fractions using a Fluorescence Activated Cell Sorting system (INFLUX 500 Cytopeia, BD Biosciences) in Lysis Buffer (50mM Tris pH=8, 10mM EDTA, 0.5% SDS, 300mM NaCl).

#### DNA extraction and sonication

DNA from sorted cells was extracted using Proteinase K treatment (200µg/ml, Thermo Scientific, EO0491) followed by phenol-chloroform extraction and sonicated to a size of 500-1,000 base pair (bp), as previously described (Hadjadj et al., 2016).

#### Immunoprecipitation using SX-8G IP-Star® Compact Automated System (Diagenode)

Immunoprecipitations from 10^5^ cells were performed using IP star robot at 4°C (indirect 200µl method, Diagenode) with an anti-BrdU antibody (10μg, purified mouse Anti-BrdU, BDg, purified mouse Anti-BrdU, BD Biosciences, #347580). Denatured DNA was incubated 5 hours with anti-BrdU antibodies in IP buffer (10mM Tris pH=8, 1mM EDTA, 150mM NaCl, 0.5% Triton X-100, 7mM NaOH) followed by 5 hours incubation with Dynabeads Protein G (Invitrogen, 10004D) (Hadjadj et al., 2016). Beads were then washed with Wash Buffer (20mM Tris pH=8, 2mM EDTA, 250mM NaCl, 1% Triton X-100). Reversion was performed at 37°C during 2 hours with a solution containing 1% SDS and 0.5mg Proteinase K followed, after the beads removal, by an incubation at 65°C during 6 hours in the same solution.

#### DNA purification and quantitative PCR

Immunoprecipitated BrdU labeled DNA fragments were extracted with phenol-chloroform and precipitated with cold ethanol. Control quantitative PCRs (qPCRs) were performed using oligonucleotides specific of mitochondrial DNA, early (*BMP1* gene) or late (*DPPA2* gene) replicating regions (Ryba et al., 2011; Hadjadj et al., 2016).

#### Amplification

Whole genome amplification was performed using SeqPlex™ Enhanced DNA Amplification kit as described by the manufacturer (Sigma-Aldrich, SEQXE). Amplified DNA was purified using PCR purification product kit as described by the manufacturer (Macherey-Nagel, 740609.50). DNA amount was measured using a Nanodrop. Quantitative PCRs using the oligonucleotides described above were performed to check whether the ratio between early and late replication regions was still maintained after amplification.

#### Universal Linkage System (ULS™) labeling, chip loading and scanning for Repli-chip experiment (microarray)

Early and late nascent DNA fractions were labelled with Cy3-ULS and Cy5-ULS, respectively, using the ULS arrayCGH labeling Kit (Kreatech, EA-005).

Same amounts of early and late-labeled DNA were loaded on mouse or human DNA microarrays (SurePrint G3 Human CGH arrays, Agilent Technologies, G4449A). Hybridization was performed as previously described (Hadjadj et al., 2016). The following day microarrays were scanned using an Agilent C-scanner with Feature Extraction 9.1 software (Agilent technologies). RT datasets are available in the Gene Expression Omnibus (GEO) database under the accession numbers GSM2111308 for U2OS, GSM2111313 for K562 and GSM2111310 for HEK293T cell lines.

### START-R suite

START-R Analyzer and START-R Viewer are open source web-based applications (doi: 10.5281/zenodo.3243339), developed using the Shiny R package (Chang et al.. 2018). START-R suite was concatenated using Docker (Supplemental Fig. S1) in order to install, use and shareit easily. START-R softwares can be used with different operating systems: Windows, Mac OS and Linux. The source code and the installation procedure are available on GitHub (https://github.com/thomasdenecker/START-R) with informations on how to use the software.

To install START-R, users should read and follow the Readme file available on GitHub web site (https://github.com/thomasdenecker/START-R/blob/master/README.md). Briefly, users should install Docker and then follow installation procedures described for each OS in the Readme file. Finally, in order to run the START-R suite, the user should double-click on the START-R file (Windows) or launch the command line (Linux / MacOS X), followed by opening an internet browser at the following URLs: http://localhost:3838/ for START-R Analyzer and http://localhost:3839/ for START-R Viewer (Supplemental Fig. S1).

### Early/Late Repli-seq data and conversion

In order to validate our softwares, we used data from the GEO database, with the accession numbers GSM2496038 and GSM2496039, corresponding to Repli-seq 46C mouse cells - Early S fraction or Late S fraction, respectively. Data were managed using different tools from GALAXY server (Afgan et al., 2018) but data can also be processed manually using specific command lines of the algorithms used on a local computer. Read mapping was obtained using Bowtie2 (2.3.4.2 version) with the very sensitive end-to-end option. Then, PCR duplicates were removed by RmDUP from SAMTools (2.0.1 version). We used BamCoverage (3.1.2.0.0 version with default parameters) with a bin size of 10kb (corresponding to the spacing of probes on microarrays) and a Reads Per Kilobase Million (RPKM) normalisation to generate a bedGraph file. A headline was added to the file to name the 4 columns (chr, start, end, gProcessedSignal for early file or rProcessedSignal for late file, respectively). Then, a script to convert and merge the bedGraph files from early and late samples to a format compatible with START-R analyzer was developed. This script is available on GitHub (https://github.com/thomasdenecker/START-R) in “supplement script” file as convert_bamcoverage_file.R.

### Validation of START-R suite using microarray data from other laboratories

We compared microarray data with Repli-seq data obtained with mouse ES46C cell line (GEO accession numbers: GSM2496037 and GSM2496038-039, respectively). Then, we analyzed with the START-R suite the microarrays data obtained by Hiratani *et al.* (2008) of D3esc and D3npc9 cell lines during mouse cells differentiation (GEO accession numbers: GSM450273 and GSM450285, respectively). As data extracted from the Nimblegen platform are in PAIR format, we used a script to convert data into a valid format for START-R Analyzer (convertPair.R, available on GitHub https://github.com/thomasdenecker/START-R in “supplement script” file).

### Validation of START-R suite using S/G1 data from multiple species

Different laboratories analyze variations of DNA copy number between G1 and S phase cells (S/G1 ratio) to study the replication timing program with Repli-seq. We used data obtained from different organisms such as Drosophila, zebrafish and human (Armstrong et al., 2018; Siefert et al., 2017; Massey et al., 2019), to validate the START-R suite (GEO accession numbers: GSM3154888 and GSM3154890 for HWT *Drosophila melanogaster* female larvae wing disc cells in S and G1 phase, respectively; GSM2282090 for 28hpf *Danio rerio* embryos; SRX3413939-40 for HEK293T human cells in S and G1 phase, respectively). As previously, reads from G1 and S fractions are mapped with Bowtie2, then PCR duplicates are removed by RmDUP tool, and Bamcoverage is used to obtain the coverage with RPKM. Then, as above, the bedGraph files are converted to a format compatible with START-R analyzer with “convert_bamcoverage_file.R.” script.

### GC content, Long Interpersed Nuclear Elements-1 (LINE-1) and gene coverage calculation

LINE-1 elements coordinates were extracted from the University of California Santa Cruz (UCSC) Genome Browser Repeat Masker track (Smit et al., 1996-2010). Two steps are required to calculate GC content: (i) DNA sequence extraction from Early, Mid, Late, Advanced, Delayed regions coordinates with the Extract Genomic DNA tool (2.2.4 version with default parameters); (ii) calculation of GC content with the GeeCee EMBOSS tool (5.0.0 version with default parameters) present in the GALAXY website (Afgan at al., 2018). Coverage tools from GALAXY website were used to determine LINE-1 content and gene coverage. All boxplots were made using R program (3.5.1 version, R Core Team, 2019).

## Data Access

All raw data generated in this study have been submitted to the NCBI Gene Expression Omnibus (GEO; https://www.ncbi.nlm.nih.gov/geo/) under accession number GSE141122. The START-R suite and the script used to perform the analysis in this study are available on GitHub at https://github.com/thomasdenecker/START-R,

## Acknowledgments

This project was supported by La Ligue Nationale Contre le Cancer (RS16/75-108 and RS17/75-135), the Groupement des Entreprises Françaises en Lutte Contre le Cancer (GEFLUC), the Institut National du Cancer INCa-10493, the IdEx Université de Paris ANR-18-IDEX-0001 and by the generous legacy from Ms Suzanne Larzat to our group.

We thank Gaëlle Lelandais for discussions during START-R elaboration and for allowing Thomas Denecker to work on this project during his PhD work. We also acknowledge the ImagoSeine core facility of the Institut Jacques-Monod, member of the France BioImaging (ANR-10-INBS-04) supported by the Region Île-de-France (E539).

